# Soft rot pathogen *Dickeya dadantii* 3937 produces tailocins resembling the tails of *Enterobacteria* bacteriophage P2

**DOI:** 10.1101/2023.08.14.553165

**Authors:** Marcin Borowicz, Dorota M. Krzyżanowska, Magdalena Narajczyk, Marta Sobolewska, Magdalena Rajewska, Paulina Czaplewska, Katarzyna Węgrzyn, Robert Czajkowski

## Abstract

Tailocins are nanomolecular machines with bactericidal activity. They are produced by bacteria to contribute to fitness in mixed communities, and hence, they play a critical role in their ecology in a variety of habitats. Here, we characterized the new tailocin produced by *Dickeya dadantii* strain 3937, a well-characterized member of plant pathogenic Soft Rot *Pectobacteriaceae* (SRP). Tailocins induced in *D. dadantii* were ca. 166 nm long tubes surrounded by contractive sheaths with baseplates having tail fibers at one end. A 22-kb genomic cluster involved in their synthesis and having high homology to the cluster coding for the tail of the Enterobacteriophage P2 was identified. The *D. dadantii* tailocins, termed dickeyocin P2D1 (phage P2-like dickeyocin 1), were resistant to inactivation by pH (3.5 – 12), temperature (4 – 50 °C), and elevated osmolarity (NaCl concentration: 0.01 – 1 M). P2D1 could kill a variety of different *Dickeya* spp. but not any strain of *Pectobacterium* spp. tested and were not toxic to *Caenorhabditis elegans*.

**Teaser:** Tailocins are nanomolecular entities similar to syringes that are produced by various bacteria to fight other microorganisms present in the same environment.

## Introduction

Under natural conditions, bacterial species inhabit shared environments, developing spatial and temporal interspecies associations and communities having complex networks of interactions (*1, 2*). In such communities, a particular member needs to continuously compete for limited resources (i.e., scarce nutrients and limited space) with most other members of the community to gain a competitive edge (*3, 4*). Given such challenging conditions, bacteria have evolved diverse strategies to successfully coexist with both closely and distantly related microbes (*5*). Such strategies, although employing a broad range of mechanisms, can be distinguished as either 1) indirect, exploitative competition that occurs through the consumption of resources and 2) direct, interference competition, where individual cells directly kill one another, limiting their lifespan (*6*). Whereas exploitative competition depends primarily on the utilization of limited resources by a strain, thereby restricting it from the competitor, interference competition relies on producing various antimicrobial agents that aim to kill other cells (*7*). These antimicrobials include, but are not limited to, broad-spectrum antibiotics, toxins, contact-dependent inhibition, effectors transported via type VI secretion system (T6SS effectors), low molecular weight bacteriocins, and tailocins (*5, 8*). A given bacterial cell may often use several such systems to gain fitness advantages in the environment (*7*). Among the systems bacteria exploit to fight competitive microbes, tailocins are now receiving increasing attention (*9, 10*).

Tailocins are syringe-like nanomolecular entities that are evolutionary and morphologically related to bacteriophage tails, type VI secretion systems, and extracellular contractile injection systems (*10*). These particles, also known as high molecular weight bacteriocins or phage tail-like particles, are chromosomally encoded and ribosomally synthesized toxins that usually express a narrow killing range, interacting only with closely-related bacterial species that typically would occupy the same niche (*9*). These agents adsorp to the surface of susceptible cells, thereby puncturing the cell envelope, leading to depolymerization of the cell membrane and, ultimately, the death of the attacked cell (*11*).

Tailocins are classified into two distinct families: rigid and contractile (R-type) and noncontractile but flexible particles (F-type) (*12*). The R-type tailocins have feature of tails of P2 or T-even bacteriophages of the family *Myovirideae* infecting *Escherichia coli*. In contrast, the F-type tailocins resemble the flexible tails of bacteriophage lambda (λ) in the family *Siphoviridae*. Although tailocins exhibit remarkable morphological similarity to bacteriophage tails of the *Myoviridae* and *Siphoviridae* viruses, it is now believed that they have evolved independently from bacteriophages and should not be considered exclusively as domesticated prophages or phage remnants that bacteria harness for their advantage (*10*).

The production of tailocins has been demonstrated both in Gram-negative and Gram-positive bacterial species. Producing strains include both human, animal, and plant pathogens and saprophytic bacteria residing in various environments (*13–16*). Until recently, tailocins have been best characterized in *Pseudomonas* species (*17*). There are, however, reports of tailocins isolated from *Clostridioides* spp., *Serratia* spp., *Xenorhabdus* spp., *Burkholderia* spp., *Kosakonia* spp., *Budvicia* spp., *Pragia* spp., *Pectobacterium* spp. as well as from other bacteria (*18*).

The omnipresence of tailocins in phylogenetically unrelated bacterial genera suggests that these particles are important for fitness in various habitats (*10*). However, the ecological role of tailocins in the natural environment of the producing strains has received little attention, especially for plant-pathogenic bacteria residing in agricultural locations. This issue is important given the diversity of conditions such bacteria encounter in such natural settings. No comprehenisve studies have addressed tailocins produced by Soft Rot *Pectobacteriaceae* (SRP) bacteria (*19*), which, due to their complex lifestyle, have a variety of spatial and temporal interactions in varied environments (*20–22*).

Plant pathogenic SRP (consisting of *Pectobacterium* spp., *Dickeya* spp., and *Musicola* spp., formerly characterized as pectinolytic *Erwinia* spp.) are a useful model for studying the environmental role of tailocins. SRP bacteria are widespread in various ecological niches, including rain and surface water, natural and agricultural bulk and rhizosphere soils, sewage, the exterior and interior of host and non-host plants as well as the surface and interior of insects (*19*). Because of the diverse environments in which SRP bacteria may be found, these pathogens may encounter various other bacteria with whom they must effectively compete.

This study aimed to assess the presence and activity of tailocins induced and isolated from pectinolytic *Dickeya dadantii* strain 3937 (*23*). This strain (formerly *Erwinia chrysanthemi* and *Pectobacterium chrysanthemi* (*24*)) is a well-known necrotrophic plant pathogen that causes soft rot disease in a variety of crop, ornamental, and other nonfood plants worldwide, causing lossess in agriculture (*25*). Strain 3937 has been widely used as a potent model system for research on the molecular biology and pathogenicity of bacteria belonging to Soft Rot *Pectobacteriaceae* for several decades (*26, 27*). While this strain continues to be the most studied strain of all *Dickeya* species its production of tailocins has not been previously described. Here, we characterize for the first time the tailocin produced by *D. dadantii* strain 3937.

## Results

### D. dadantii 3937 produces tailocins

Cells of *D. dadantii* 3937 treated with mitomycin C produced syringe-like macromolecular structures resembling bacteriophage tails. Following convention, the tailocins produced by *D. dadantii* were named dickeyocins. Imaging using TEM and AFM revealed that these structures consist of a central rod-shaped core (tube) wrapped in a contractable sheath (Fig. 1). Both the tube and the sheath were built of multiple protein subunits, with a clearly visible helical arrangement of the subunits in the sheath. The average length of the dickeyocins produced by strain 3937 was 166±7 nm. When the sheath entirely covered the tube (in the extended, ‘loaded’ form), the individual dickeyocin has a diameter of 23±2 nm. When the sheath contracts, it revealed an internal tube of a length of 92±7 nm with an attached spike at the proximal end. Fibers were visible at the distal end of the sheath (Fig. 1).

**Fig. 1.**
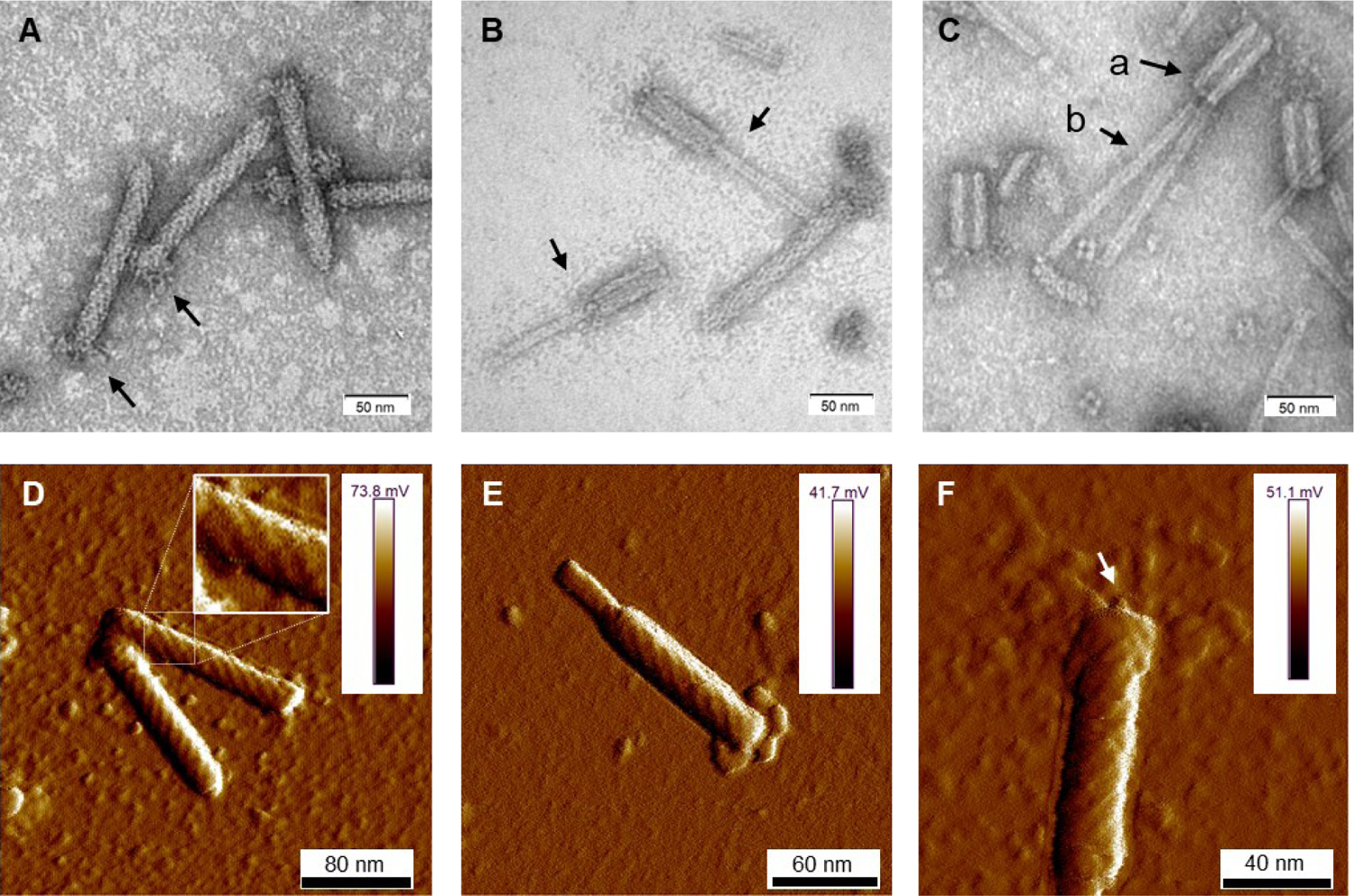
The morphology of tailocins from *D. dadantii* 3937. Images were obtained using TEM (A-C) and AFM (D-F). The individual panels show: (**A**) particles in an extended (active) form. Arrows indicate the positioning of fibers; (**B**) contracted particles indicated by arrows; (**C**) tailocins disintegrated by heat treatment (50 ⁰C); Arrows indicate: **a** – empty sheath with a hollow inner channel, **b** – tube (core) separated from the sheath; (**D**) two extended particles. A fragment of the photo was magnified to display the helical arrangement of protein subunits in the sheath; (**E**) a contracted molecule in an AFM image; (**F**) an extended particle with an arrow pointing at the presumed tube-attached spike.

The yield of tailocins purified from mitomycin-induced cells of *D. dadantii* 3937 equaled approx. 10^11^ particles mL^-1^ of culture. The particles could also be isolated from non-treated cells, indicating a low basal production level. However, induction with mitomycin C increased the yield approx. 10-100 fold. We employed three different methods to estimate the concentration of phage-tail-like particles in the tested preparations. The results were consistent between these methods, as well as for the independently-obtained batches of purified dickeyocins (Table S2). The average concentration from three independently obtained samples was 10^6^ relative units (AU) mL^-1^, 10^11^ killing particles mL^-1,^ and 10^11^ particles mL^-1^, according to the spot test, the Poisson distribution killing method, and the NanoSight measurements, respectively. Furthermore, comparing the results from the activity-based Poisson method and the direct particle count indicated that most phage-like particles purified from the cultures of *D. dadantii* strain 3937 were undamaged and in the extended (’loaded’) form. This was in line with the microscopic observations done with TEM and AFM.

### Dickeyocins from D. dadantii strain 3937 are phylogenetically related to the tail of P2 bacteriophage

Proteins in the dickeyocins were separated by SDS-PAGE and sequenced. Eight cleary distinguishable bands were excised from the gel, and the digested peptides were analyzed by MS. The peptides could be readily mapped to six *D. dadantii* proteins with annotations implying their phage relationship: phage baseplate assembly protein (encoded in locus Dda3937_00029 / DDA3937_RS12055), baseplate assembly protein J (Dda3937_00030 / DDA3937_RS12060), tail fiber protein (Dda3937_04606 / DDA3937_RS12070), putative side tail fiber protein (Dda3937_03808 / DDA3937_RS12100), major sheath protein (Dda3937_03810 / DDA3937_RS12110), and major tail tube protein (Dda3937_03811 / DDA3937_RS12115) (Fig. 2). Genes encoding these six proteins were mapped to a single ca. 22-kb region in the genome of *D. dadantii* strain 3937 (GenBank accession number: NC_014500.1: genome location: 2,734,508 to 2,757,061) (Fig. 2C). This genomic region contained 28 genes, from which 16 genes encoded various bacteriophage structural proteins.

**Fig. 2.**
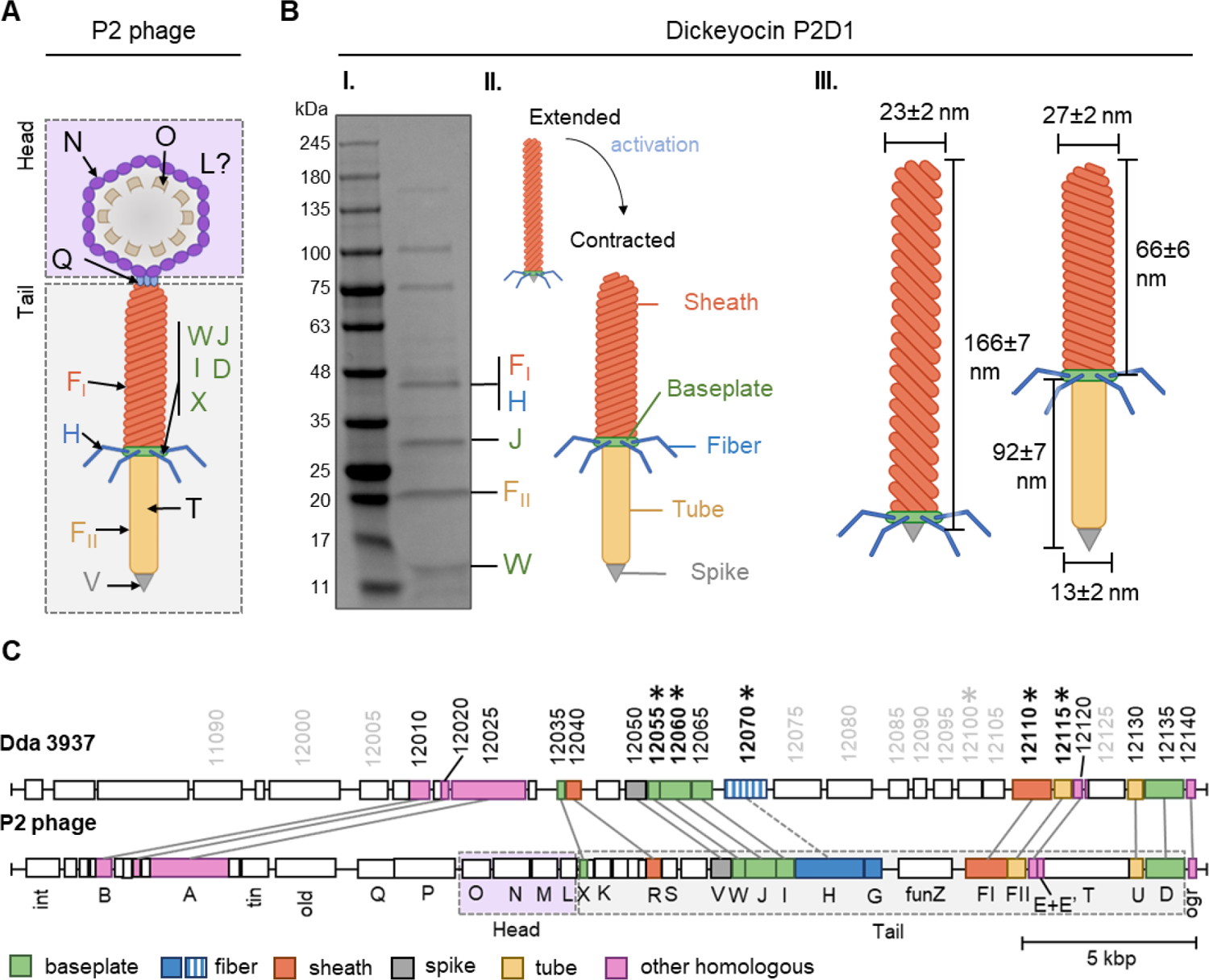
The building blocks of tailocins of *D. dadantii* strain 3937 and their encoding genes in relation to those of the *Enterobacteria* P2 bacteriophage. **Panel A** shows a schematic representation of the structure of the P2 phage, together with the nomenclature of the building proteins. The graph was prepared based on Cristi and Calendar, 2016 (PMID: 27144088, https://pubmed.ncbi.nlm.nih.gov/27144088/). WJIDX – baseplate proteins, F_I_ – sheet, H - fiber, F_II_ – tube, V – spike, T – tape measure protein, Q – portal protein, N – major capsid protein, O – capsid scaffold, L – head completion protein. **Panel B** shows: **I.** an SDS-PAGE separation of proteins from mitomycin-induced cultures of *D. dadantii* 3937. Bands containing proteins homologous to those of the P2 phage are provided with the respective protein designations; **II.** schematic representation of extended and contracted forms of dickeyocin P2D1; **III.** dimentions of extended and contracted forms of dickeyocin P2D1. **Panel C** shows the alignment between the complete genome sequence of the P2 phage (NC_001895; 33 593 bp) and a region of the same length in the genome of *D. dadantii* 3937 (NC_014500.1; range: 2723487 to 2757061). The numbering of genes in *D. dadantii* corresponds to the numbering of loci in the genome (locus tag prefix DDA3937_RS). Homologous proteins are marked with the same color.

The nucleotide sequence of the 22-kb putative dickeyocin-encoding fragment in *D. dadantii* strain 3937 showed the highest similarity to a prophage region in *D. dadantii* strain XJ12 (100% query coverage (qc), 99.35% identity). Highly similar regions were also found in the genomes of several other strains of *D. dadantii*, as well as *D. solani*, *D. dianthicola*, and *D. fangzhongdai* (Data S1). Strains of *D. zeae* showed lower similarity, with query coverage between 54-76% and identity of 83-84%. A much lower score was observed for the next best hit – *Musicola paradisiaca* (formerly *Dickeya paradisiaca*). Other non-*Dickeya* microorganisms with regions showing some degree of homology included *Serratia* sp. ATCC 39006 (Data S1) (with 10% query coverage and 81.45% identity).

Importantly, at the nucleotide level, the dickeyocin region of *D. dadatii* strain 3937 showed no significant similarity to the known carotovorocin Er cluster of *P. carotovorum* Er (Genbank accession: AB045036) (*28*). Moreover, the amino acid sequence of the sheath protein of tailocin from strain 3937 showed only a 34% identity to that of carotovorocin Er, with a query coverage of 83%.

The ca. 22-kb cluster encoding dickeyocin from *D. dadantii* strain 3937 was also surveyed against the collection of viral sequences (NCBI taxid: 10239), yieldingonly low similarity scores (7% query coverage, 78% identity for the best hit).

Furthermore, in an attempt to find undomesticated phages related to dickeyocin we searched the viral protein database, using three structural proteins of dickeyocin as queries: sheath protein (WP_013318223), tail tube protein (WP_013318224), and baseplate assembly protein J (WP_013318212). Based on this search, proteins in dickeyocin exhibited homology to proteins of *Salmonella* phages SW9 and PSP3, *Erwinia* phage Etg, *Enterobacteria* phage fiAA91-ss, *Peduovirus* (bacteriophage) P2, several *Escherichia* and *Yersinia* phages, as well as to multiple poorly-characterized phages of bacteria in class *Caudoviricetes*, derived from the human metagenome (Data S1). The best-characterized phage having a high homology to dickeyocin was *Enterobacteria* phage P2 – a *Myoviridae* phage that infects *Escherichia coli* and other hosts, including *Salmonella* and *Klebsiella* (*29, 30*). For the three investigated dickeyocin proteins, their amino acid identity towards their P2 homologs ranged from 70-79%, with 100% query coverage (Data S2). Therefore, we used the *Enterobacteria* P2 bacteriophage as a reference to assign functions to proteins in dickeyocin, as well as to investigate the genetic rearrangements in the dickeyocin cluster in relation to the fully-functional phage P2 (Fig. 2C). The dickeyocin cluster in the genome of *D. didantii* strain 3937 lacked sequences associated with the phage head (proteins O, M, N, L), as well as the tape measure protein. There was also a difference in the genetic content of the intergenic region of the tail, as well as low homology of proteins building the tail fibers (31% query coverage, 54% identity). Following the present naming convention, we named the newly characterized tailocins from *D. dadantii* 3937 **dickeyocin P2D1** (<u>P2</u>-like <u>D</u>ickeyocin <u>1</u>).

### Dickeyocin P2D1 expresses bactericidal activity exclusively against members of Soft Rot Pectobacteriaceae

Fifty-two bacterial strains were surveyed for their sensitivity to tailocins induced from *D. dadantii* strain 3937. These included 41 strains of different species and subspecies of *Dickeya* and *Pectobacterium*, as well as 7 other bacterial strains belonging to the *Enterobacteriaceae* family, 3 *Pseudomonas* spp., and a single strain of *Staphylococcus aureus* representing Gram-negative bacteria (Table S1). Bactericidal activity was observed against eight strains (*D. dadantii* subsp. *dieffenbachie* strain NCPPB 2976, *D. dianthicola* strains NCPPB 3534 and IPO 980, *D. undicola* CFBP 8650, *D. zeae* strains NCPPB 3532 and 3531, *D. oryzae* strain CSL RW192, and *Musicola paradisiaca* strain NCPPB 2511 (old name: *Dickeya paradisiaca* strain NCPPB 2511)), but not against any of the *Pectobacterium* spp. tested (Table S1, Fig. 3). Likewise, dickeyocin P2D1 was inactive against any of the non-SRP strains included in the screening assay: 7 *Enterobacteriaceae* strains (*Serratia marcescens* ATCC 14756, *Citrobacter freundii* ATCC 8090, *Escherichia coli* ATCC 8739, *Escherichia coli* ATCC 25922, *Escherichia coli* OP50, *Klebsiella quasipneumoniae* ATCC 700603, *Klebsiella aerogenes* ATCC 51697), 3 *Pseudomonas* spp.: (*Pseudomonas aeruginosa* PA14, *Pseudomonas aeruginosa* PAO1, *Pseudomonas donghuensis* P482), and *S. aureus* ATCC 25923 indicating their limited range of bactericidal activity (Fig. 3).

**Fig. 3.**
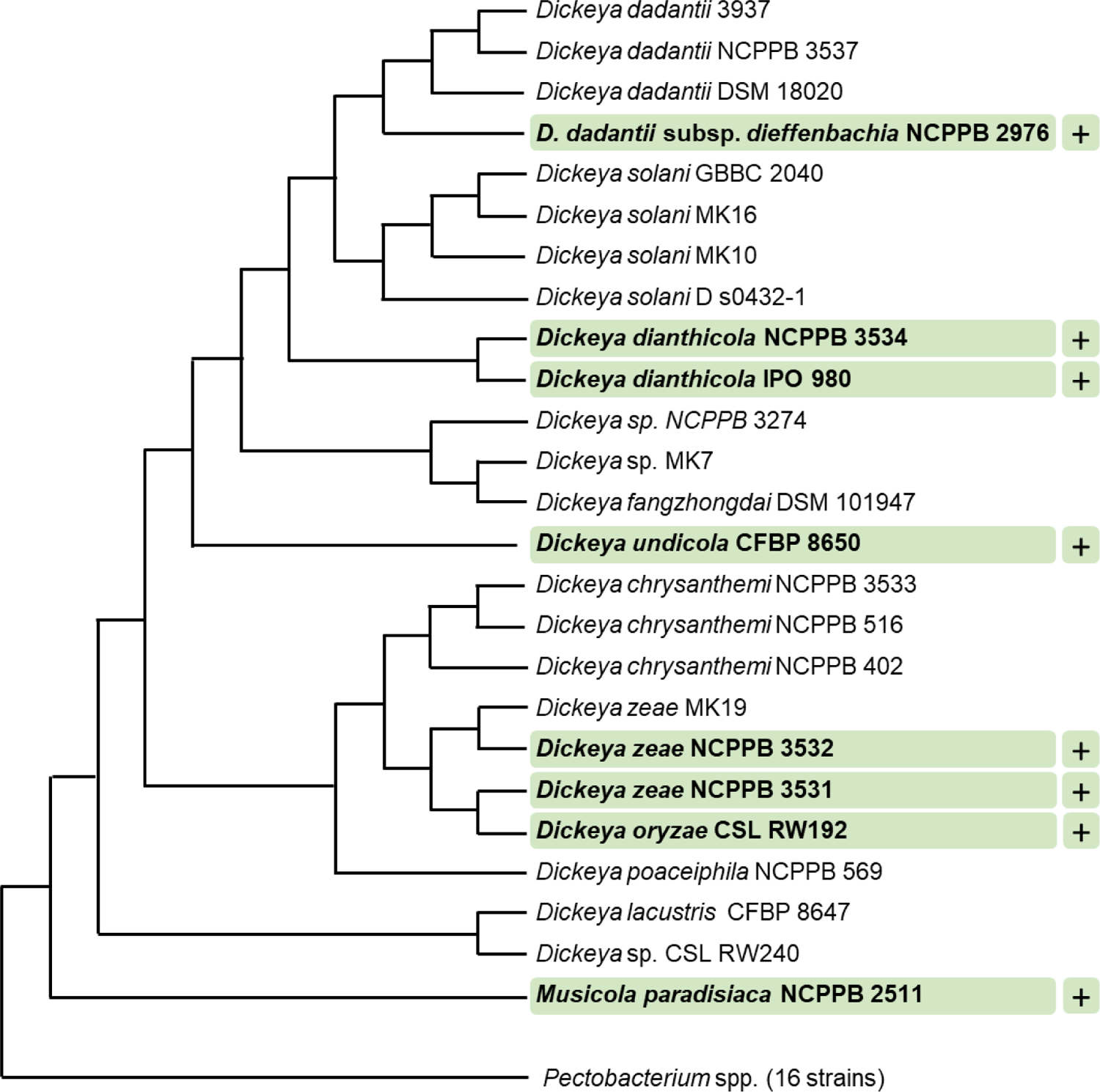
Target range of tailocins from *D. dadantii* 3937. Strains susceptible to P2D1 dickeyocin were marked with a plus (+). A phylogenetic tree for 41 SRP strains was obtained using EDGAR 3.2. It was built out of a core of 1616 genes per genome (nucleotide sequences), 66256 genes in total. The core had 1560951 bp per genome, 63998991 in total. The lengths of the branches do not reflect the phylogenetic distance. All analyzed *Pectobacterium* spp. were collapsed. This included 16 strains: *P. cacticida* CFBP 3628, *P. fontis* CFBP 8629, *P. betavasculorum* CFBP 2122, *P. peruviense* CFBP 5834, *P. atrosepticum* SCRI1043, *P. atrosepticum* NCPPB 549, *P. parmentieri* CFBP 8475, *P. parmentieri* SCC3193, *P. polonicum* DPMP 315, *P. punjabense* CFBP 8604, *P. actinidiae* LMG 26003, *P. brasiliense* LMG21371, *P. polaris* NCPPB 4611, *P. versatile* CFBP 6051, *P. carotovorum* CFBP 2046, *P. aroidearum* NCPPB 929.

### P2D1 killing efficiency is bacterial species-dependent

We determined the kinetics of killing of eight susceptible strains treated with P2D1. The share of intact cells at two time points: 20 and 120 minutes post treatment is shown in Fig. S1. In all the cases, the killing of the susceptible cells was fast; in the first 20 minutes, the most killing was observed for *D. dianthicola* strain IPO 980 (ca. 60% reduction of cell numbers). In contrast, the slowest killing caused by dickeyocin P2D1 was observed for *D. dianthicola* strain NCPP 3534 (ca. 22% reduction of cell numbers). For the other 6 strains tested, on average, a 45 to 60% reduction in cell numbers was observed after this same time (20 min.). About 50 to 60% killing of all strains by dickeyocin P2D1 was seen after 120 min-incubation (Fig. S1).

### P2D1 dickeyocin directly punctures the cell envelope of susceptible bacterial strains

We assessed the interaction of dickeyocin P2D1 on cells of the susceptible *M. paradisiaca* strain at the single-cell level using transmission electron microscopy (TEM). Images clearly showed that dickeyocin P2D1 adsorbed to the bacterial cell envelope (Fig. 4 AB), followed by puncturing the outer cell membrane (resulting from the conformational change of dickeyocin P2D1 from an extended ‘loaded’ to a contracted form) (Fig. 4 CD), creating a pore connecting the cytoplasm of the cell with the external environment and leading eventually to the death of the attacked cell.

**Fig. 4.**
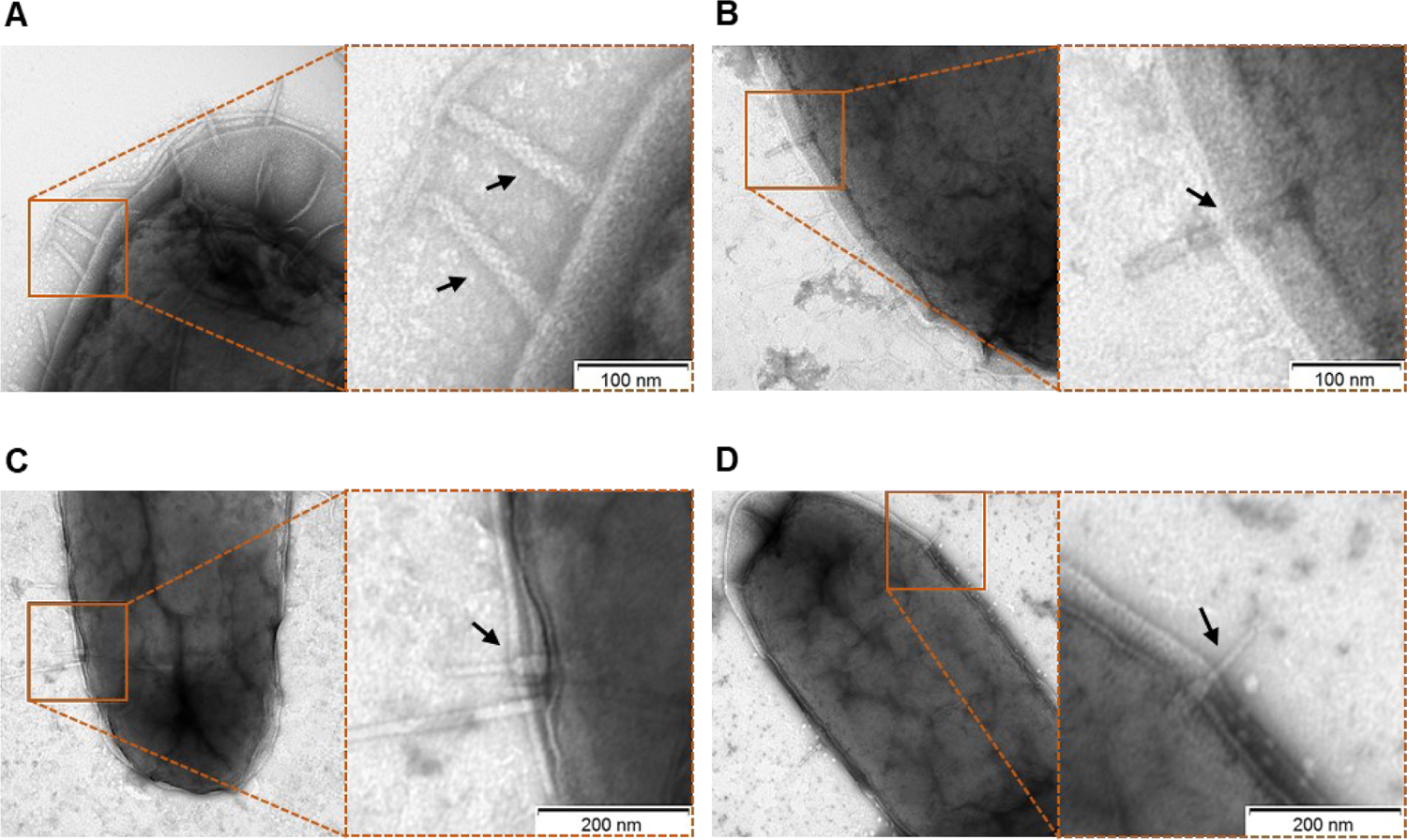
Interaction of P2D1 dickeyocins with the cells of a susceptible strain *M. paradisiaca* NCPPB strain 2511, evaluated using transmission electron microscopy (TEM). Bacterial cells of *M. paradisiaca* NCPPB 2511 from overnight culture in TSB were harvested and washed with PBS. Such prepared cells were incubated with PEG-purified dickeyocins (approximate concentration in bacterial suspension 10^6^ particles mL^-1^) for 20 min, followed by obtaining images with TEM. For all panels, the TEM images on the right are the enlarged sections of the original micrographs on the left. Arrows indicate P2D1 attached to the surface of bacterial cells, either in extended (A-B) or contracted (C-D) form.

### Dickeyocin P2D1 is able to bind to nonviable (dead) bacterial cells

A cell adsorption assay was employed to investigate whether the dickeyocin P2D1 can adsorb to nonviable (dead) and viable bacterial cells of the susceptible or resistant bacterial species. Dickeyocin P2D1 was incubated either with viable or chloramphenicol-killed cells of susceptible *M. paradisiaca* or with viable or antibiotic-killed cells of resistant *D. dadantii* for 40 min and the remaining dickeyocin P2D1 in the medium was measured. Incubation of dickeyocin P2D1 both with the dead and viable cells of *M. paradisiaca* resulted in the total loss of the tailocin activity, indicating that it bound equally efficiently to viable as well as nonviable cells of the susceptible bacterium. No adsorption of dickeyocin P2D1 to either dead or alive cells of the resistant *D. dadantii* cells was observed (Fig. S2).

### Production of dickeyocin P2D1 is not associated with T6SS

To determine whether dickeyocin P2D1 production in *D. dadantii* depends on the type VI secretion system (T6SS), dickeyocin P2D1 production was induced in *D. dadantii* mutant A5587 (*31*) that carries an insertional mutation in the *tssK* gene (type VI secretion system baseplate subunit TssK) – an essential subunit of the type VI secretion apparatus (*32*). The tailocins obtained from the *D. dadantii* T6SS mutant using standard mitomycin induction methods were morphologically indistinguishable and similarly abundant compared to dickeyocin P2D1induced in the wild-type *D. dadantii* strain 3937 (Fig. S3). These results suggest that at least *tssK* gene, essential for the proper function of T6SS, does not affect dickeyocin P2D1 production, morphology, or release from the cell.

### Activity of P2D1 dickeyocins is modulated by environmental conditions and enzyme treatments

Dickeyocin P2D1 was stable at temperatures between 4 and 42 °C, showing no significant loss in activity following 24 h incubation at these temperatures. Temperatures above 50 °C, however, led to a 32-fold loss in activity while incubation at 65 and 80 °C (Fig. 5 A) led to a complete loss of activity. Likewise, a single freeze-thaw cycle negatively affected the stability of dickeyocin P2D1. Freezing at −20 °C resulted in a 128-fold reduction in activity, while no bactericidal effect remained after freezing at −80 °C (Fig. 5 A). We tested the stability of P2D1 under 7 different pH conditions (2, 3.5, 5, 7, 9, 10.5, and 12). The particles remained active after 24 h incubation in pHs ranging from 3.5 to 12. However, their bactericidal properties were lost at pH 2 (Fig. 5 B). To determine whether dickeyocin P2D1 that was inactivated at low pH could be restored to an active (extended, ‘loaded’) form, we diluted the pH 2-treated, inactive dickeyocin P2D1 in PBS buffered to pH 12 to achieve a neutral pH environment.

**Fig. 5.**
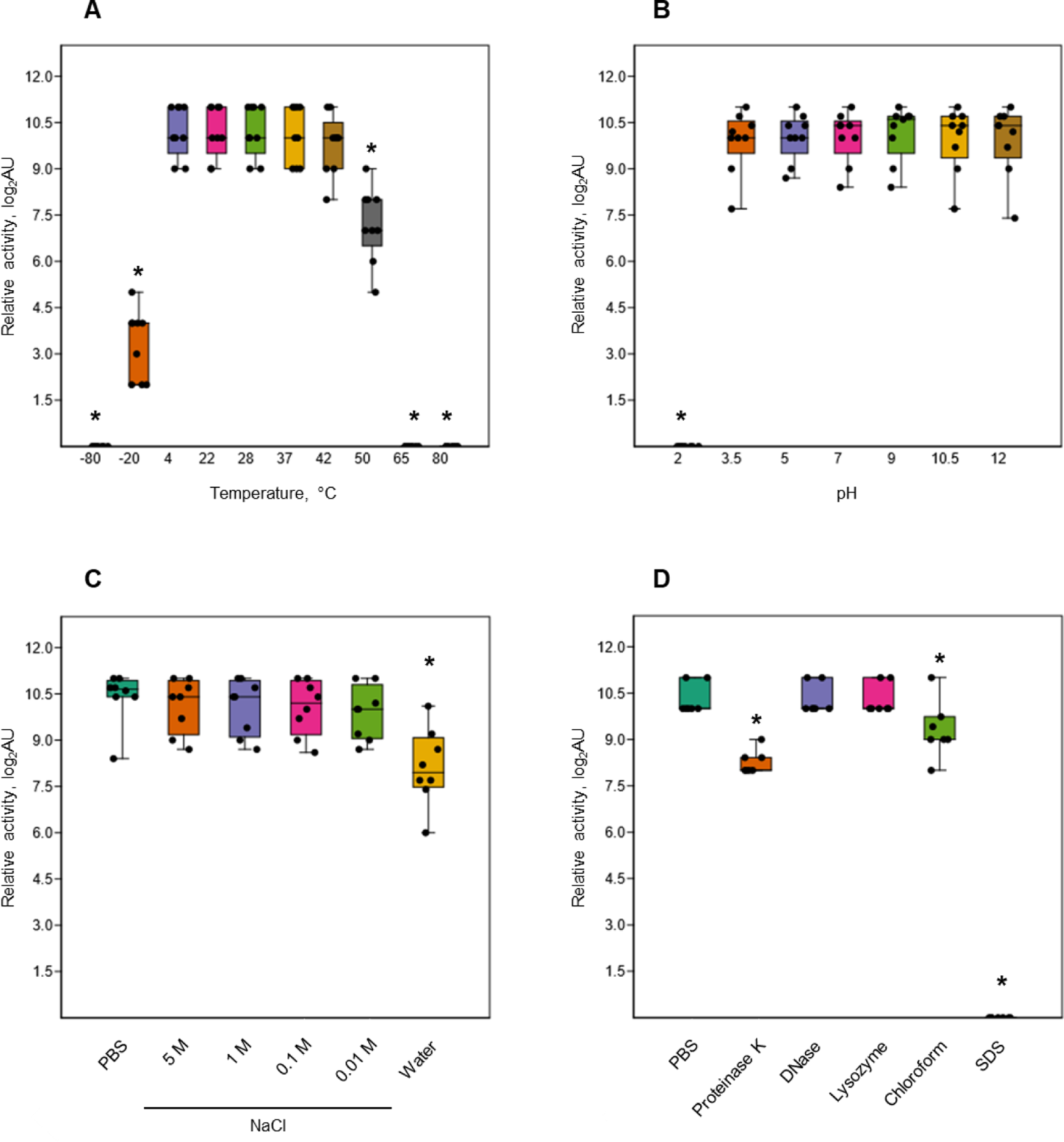
Stability of P2D1 dickeyocins from *Dickeya dadantii* strain 3937 upon different treatments. The graphs show the influence of temperature (A), pH (B), salinity (C), as well as enzymatic digestion (proteinase K, DNase, and lysozyme), chloroform treatment, and SDS detergent (D) on bactericidal activity of P2D1 against *M. paradisiaca* NCPPB 2511. For temperature, pH/osmolarity, and enzyme/detergent/organic solvent treatments, dickeyocins stored in PBS pH 7, were used as a control. The results are shown as a box plot; whiskers reflect the maximum and minimum, box sides reflect the first and third quartile and the bars reflect medians. Points indicate particular measurements (n=9). The asterisks (*) represent statistically significant differences (p<0.05) between the treatment and the control using the Mann-Whitney test.

Dickeyocin P2D1 did not regain bactericidal activity at pH 7. Osmotic conditions generated by NaCl added to water at concentrations between 0.01 to 5 M had no adverse effect on the activity of dickeyocin P2D1 within 24 hours when compared to controls (dickeyocin P2D1 in PBS containing 0.137 M NaCl and 0.0027 mM KCl). In contrast, incubation of dickeyocin P2D1 in deionized water led to a 4-fold loss in activity (Fig. 5 C). Dickeyocin P2D1 activity was significantly (p<0.05) reduced by treatment with proteinase K and chloroform for 1 hour (4-fold and 2-fold loss of activity, respectively). Complete loss of activity was seen after incubation in 1% SDS. In contrast, neither lysozyme nor DNase influenced dickeyocin P2D1 activity (Fig. 5 D).

### Dickeyocin P2D1 is nontoxic for Caenorhabditis elegans

No inhibition of the survival of *C. elegans* was seen after its exposure to high concentraiotns of purified dickeyocin P2D1 (Fig. S4). The average survival rate of *C. elegans* cultivated in the presence of P2D1 dickeyocins was ca. 98-99% and was not significantly statistically different from that of nematodes grown without dickeyocin P2D1. The survival of *C. elegans* was not dickeyocin P2D1-concentration-dependent, as similar survival rates were recorded for all dickeyocin concentrations tested (Fig. S4). Furthermore, dickeyocin P2D1 did not exhibit bactericidal activity against *E. coli* OP50 – the food source for *C. elegans*.

## Discussion

Our understanding of the mechanisms underlying bacterial competitive behaviors remains limited. Although numerous bacterial species are known to produce and exploit tailocins for their competitive advantage, little is known of how commonly such entities are employed by plant-pathogenic bacteria, including pectinolytic, necrotrophic SRP pathogens, to enable their environmental fitness (*9, 10*). This study revealed the presence of tailocins in one of the broadest characterized and economically important member of the Soft Rot *Pectobacteriaceae* family – strain *D. dadantii* 3937. We identified and characterized in detail a novel phage tail-like particle – dickeyocin P2D1, produced by this strain not only upon mitomycin C treatment, but also constitutively in culture. Although tailocins were initially noted in a limited number of *Dickeya* spp. (strains of *Erwinia chrysanthem* dissimilar from strain 3937) isolated from several crop and ornamental plants in the past (1960s and 1970s) (*33*), these particles were not characterized in detail, probably due to the lack of appropriate molecular techniques at that time.

Dickeyocin P2D1 has a typical morphology and expresses features similar to those of other R-type tailocins described so far for a large group of bacteria, including *Escherichia coli* (*34*), *Burkholderia cenocepacia* (*35*), *Pseudomonas aeruginosa* (*10*), *Proteus vulgaris* (*36*) and *Yersinia enterocolitica* (*37*). Likewise, tailocins were also identified in *Pectobacterium* spp. (*28, 38, 39*) – plant pathogens closely related to *Dickeya* spp. The tailocin of SRP bacteria best characterized to date is carotovorocin Er isolated from *P. carotovorum* Er (*39, 40*). Interestingly, as shown in this study, despite the morphological similarities of dickeyocin P2D1 and carotovorocin Er and the rather close phylogenetic relationship and the lifestyle shared by the producer strains, these tailocins show little sequence homology. This suggests that dickeyocin P2D1 and carotovorocin Er originated from different phage ancestors and that the ability to produce tailocins was acquired several times and independently by various members of the SRP family.

The cluster encoding dickeyocin P2D1 exhibits high homology with that encoding the tail of *Enterobacteria* phage P2, indicating that these two probably had a common ancestor. Phage P2 is a temperate bacteriophage belonging to the *Myoviridae* family, commonly found in genomes of strains belonging to the *Proteobacteria* phylum (*30*). Its presence has been noted in at least 127 genera in 32 *Proteobacteria* families (*41*), indicating its extreme ubiquitousness in the bacterial world. Accordingly, R-type tailocins resembling the tail of phage P2 have been extensively studied and are now the best characterized to date (*5, 10*). To explore the occurrence of R-type tailocins in SRP, we searched available *Dickeya* spp. and *Pectobacterium* spp. genomes for regions carrying phage tail-like genes homologous to those encoding dickeyocin P2D1 and having organizational similarity. Our analysis revealed the presence of P2D1-like clusters in several *Dickeya* spp. strains, including *D. solani*, *D. dianthicola, D. zeae*, and *D. fangzhongdai*. The P2D1-like clusters were, however, absent in all *Pectobacterium* spp. genomes as well as in genomes of the other bacterial species unrelated to SRP analyzed in our study. The lack of the dickeyocin P2D1 cluster both in *Pectobacterium* spp. and in bacteria phylogenetically distant to SRP may indicate a strong and sustained phylogenetic association between dickeyocin P2D1and *Dickeya* spp. This is somewhat unexpected given that various *Dickeya* and *Pectobacterium* are often present in the same environment (e.g., soil, water, plant surface) (*19*). Under such conditions, horizontal gene transfer would frequently occur between these phylogenetically related SRP species (*42*).

The restricted occurence of genes encoding dickeyocin P2D1 in the genomes of *Dickeya* spp. aligns with its narrow bactericidal activity. In our study, dickeyocin P2D1 exclusively targeted *Dickeya* species as well as *Musicola paradisiaca* (former *Dickeya paradisiaca*) but not any other bacteria tested. It is noteworthy, however, that these *Dickeya* strains differed somewhat in the suspectibiltiy to dickeyocin P2D1 while these differences were unrelated to the phylogenetical distance between them. This is in line with other studies (*43–45*) that showed that the host range of pyocins, tailocins produced by *Pseudomonas* spp., is usually restricted to kin strains (*9*). Likewise the tailocins of *Escherichia coli* (*46*) and *Yersinia enterocolitica* (*37*) also killed only related taxa. Contrarily, some recent reports have revealed tailocins that target different species of the same genus and/oreven different genera. For example, *Burkholderia cenocepacia* tailocins were active against other *Burkholderia* species (*35*) and those of *P. fluorescens* were active against *Xanthomonas vesicatoria* (*40*). The quite narrow target range of dickeyocin P2D1 is somewhat surprising given that multiple species of SRP bacteria are often found together in the same infected plant (*47–49*). In such a setting these various species would be expected to experience intense competition and would be expected to benefit from a promiscuous tailocin (*50*). It thus appears that in such mixed infections, *Dickeya* spp. and *Pectobacterium* spp. compete using mechanisms unrelated to phage-tail-like particles, as reported previously (*51, 52*), but this conjecture requires more experimental support.

Our study revealed that dickeyocin P2D1 is produced even without induction with mitomycin C, although such induction boosts the tailocin production 10 to 100-fold. To precisely estimate the concentration of P2D1 particles, we used a well-established Poisson distribution killing method (*53, 54*) as well as a new approach we developed. The Poisson distribution killing method is an indirect enumeration method as it employs the number of survivors in tailocin-treated bacterial suspensions. Advantage of indirect methods is that they provide the number of active (’loaded’) tailocin particles in a sample. In this study, we have complemented the indirect counting approach by direct enumeration of tailocin particles with NanoSight NS300. Direct particle counts avoid the need for culturing, and when combined with killing-based methods, enables an estimation of the fraction of active tailocins in a preparation. This appears to be a novel application of NanoSight to evaluate the quality and the active fraction of tailocin preparations.

One of the immediate applications of dickeyocin P2D1 would be in controlling SRP infections in crops (*18*). A major practical limitation of tailocins as therapudic agents is their narrow bactericidal range. This can, however, be at least partially overcome by using cocktails of tailocins having different bactericidal ranges, similarly as in the use of bacteriophages (*55*). Despite this limitation, tailocins have shown to be effective antibacterial agents in various applications (*56, 57*). They have several benefits compared to other biological control agents evaluated to date (*18*). One of the biggest advantages of tailocins is their incapacity to proliferate on-site after application. Likewise, tailocins themselves cannot spread via transduction or transformation mechanisms because only the protein products and not the encoding nucleic acids are employed. The high target selectivity of tailocins also prevents disruption of sother, often beneficial bacteria present in the same niche. Likewise, since dickeyocin P2D1 was nontoxic to *C. elegans*, a model eukaryotic organism, such tailocins are unlikely to impact other eukaryotic organisms present in soil and/or on plants (*58*). Additional studies are needed to fully explore the potential of dickeyocin P2D1 to achieve plant disease control, such as targeting production, the long-term effectiveness, and consistency of control under field conditions, including stability, formulation, and eco-toxicological risks. Studies of the *in planta* expression of dickeyocin P2D1 should provide great insight into the interactions of *Dickeya* species with each other and with other bacteria under natural and agricultural conditions.

## Materials and Methods

### Bacterial strains and culture conditions

All bacterial strains included in this study are listed in Supplementary Table S1. Bacteria were routinely propagated in liquid Trypticase Soy Broth (TSB, Oxoid) or on solid Trypticase Soy Agar (TSA, Oxoid) at 28 °C for 24 h. Liquid cultures were agitated during incubation (120 rpm). When required, bacterial cultures were supplemented with kanamycin (Sigma-Aldrich) to a final concentration of 50 µg mL^-1^.

### Induction, purification, and concentration of tailocins from *Dickeya dadantii* 3937 culture

*D. dadantii* strain 3937 was grown overnight (ca. 16 h) in TSB at 28 °C with shaking (120 rpm). The cultures were then rejuvenated by diluting them 1:40 in 250 mL of fresh TSB medium. The diluted culture grew for 2.5 hours under the same conditions. Such prepared 3937 cultures were then supplemented with mitomycin C (Abcam, Poland) to a final concentration of 0.1 µg mL^-1^ to induce the production of tailocins. Following mitomycin C treatment, the cultures were incubated for another 24 h at 28 °C with shaking (120 rpm). Bacterial cells were then removed by centrifugation (10 min, 8 000 RCF), and the supernatant containing putative particles was collected and filtered through a sterile 0.2 µm PES (polyether sulfone) membrane filter using the Nalgene Rapid-Flow Sterile Disposable Filter Units (Thermo Fisher Scientific). Finally, to precipitate tailocins, PEG-8000 (Promega) was added to the filtrate to a final concentration of 10%, and the sample was incubated at 4°C on a magnetic stirrer for the next 16-24 h. The tailocins were collected by centrifugation (1 h, 16 000 RCF, 4 °C) and resuspended in 5 mL of Phosphate Buffered Saline (PBS, pH 7.2, Sigma-Aldrich). The resulting pellet was resuspended in 1/50 of the initial volume of the initial sample. The purified particles were stored at 4 °C.

### Initial qualitative screen of tailocins for the bactericidal activity

The activity of the purified and concentrated particles was initially tested qualitatively on a limited panel of bacterial strains (=17 strains, Supplementary Table S1) using a spot test assay as described before (*35, 59*).

### Microscopic imaging

Tailocins were imaged using both transmission electron microscopy (TEM) and atomic force microscopy (AFM), as described earlier (*60*). TEM analyses were done at the Laboratory of Electron Microscopy (Faculty of Biology, University of Gdansk, Poland). For TEM analysis, particles obtained as described above were adsorbed onto carbon-coated grids (GF Microsystems), stained with 1.5% uranyl acetate (Sigma-Aldrich), and directly visualized with an electron microscope (Tecnai Spirit BioTWIN, FEI) using a previously described protocol (61). At least ten images were taken for each preparation to estimate the diameters of the particles. For the AFM analysis, purified and PEG-concentrated particles were used directly without further preparations. AFM imaging was conducted in air mode using the Bioscope Resolve microscope (Bruker), in ScanAsyst (Peak Force Tapping) mode, employing the SCANASYST-AIR probes (f0 7.0 kHz, diameter <12 nm, k: 0.4 N/m) as described earlier (62). Similarly, as described for TEM analysis, for AFM, at least ten images were taken for each preparation to estimate the diameters of the particles.

### Determination of the concentration of tailocins

To estimate the concentration of tailocins, three independent methods were applied.

### Direct particle count with NanoSight NS300

NanoSight NS300 instrument (Malvern Panalytical), equipped with an sCMOS camera and a Blue488 laser, was used to directly assess the concentration and size distribution of particles obtained after induction and purification, as described above. The tailocin samples were diluted 1000 times in sterile PBS buffer pH 7.2 (Sigma-Aldrich) to acheive the optimal concentration for observation. The camera gain was set to 14, the number of captures per sample equaled 5, each lasting 60 s, and the detection threshold was set to 5 as suggested by the manufacturer. Measurements were conducted at room temperature (ca. 22-23 °C). Three biological replicates were used to determine the concentration and size distribution of the obtained tailocins, and the results were averaged for further analysis.

### Semiquantitative estimation by a spot test (*35*)

Samples interrogated for tailocins were serially 2-fold diluted in PBS pH 7.2 (Sigma-Aldrich). Two µl of each dilution were spotted onto TSA plates overlayed with 15 ml of soft top agar (Nutrient Broth (NB, Oxoid) with 7 g L^-1^ agar). Before pouring, the soft top agar was cooled to 45 °C and inoculated with 250 µl of an overnight culture of a tailocin indicator strain (susceptible strain *M. paradisiaca*). Plates were incubated overnight at 28 °C. The highest dilutions of particles capable of cell lysis, visible as plaques (halos) in the bacterial lawn in soft top agar, were determined. Each tailocin dilution was tested in triplicates and the entire experiment was repeated two times with the same setup. The reciprocal of the highest dilution causing a visible plaque was defined as the value of the relative activity in arbitrary units (= 1 AU).

### Poisson distribution killing method (*35*)

Tailocins were also quantified using the Poisson distribution killing method based on the protocol described by (*35*) and initially introduced by others (*53, 54*). This method is based on the number of bacteriocidal events, determined from the number of bacterial survivors in a population with a known number of initial viable cells. To determine this, ten µl of undiluted and 10-fold diluted samples of tailocins were added to 100 µL of an overnight TSB (Oxoid) culture of a susceptible strain *Musicola paradisiaca* NCPPB 2511 (10^8^ CFU mL^-1^) and incubated for 40 min at 28 °C with shaking (120 rpm). As a negative control, PBS pH 7.2 was used instead of the tailocins suspension. Each combination was tested in triplicates. After incubation, the suspensions were serially diluted up to 10^-7^ in PBS and plated in triplicate on TSA plates. The colonies that emerged following overnight incubation at 28 °C were enumerated to calculate the bacterial survival ratio (*S*) in the treated samples compared to the negative control. S was calculated as the number of viable bacteria in a sample incubated with tailocins divided by the number of viable bacteria in the negative control. The number of lethal events per bacterial cell (*m*) was calculated as *m* = -ln(*S*). The total number of active killing particles per milliliter (based on the assumption that tailocins adsorption to bacterial cells in each sample was quantitative within the first 40 min incubation period) was calculated by multiplying *m* by the initial number of bacterial cells per milliliter (CFU mL^-1^).

### Target range of the tailocins isolated from *D. dadantii* 3937

Overnight cultures of the investigated strains were prepared in 1 mL aliquots of TSB in 2 mL microcentrifuge tubes (Eppendorf) which were incubated horizontally with shaking (120 rpm) at 28 °C for 16-24 h. The overnight cultures were used as an inoculum in a spot test carried out on 48-well plates (Greiner). Briefly, 10 µL of the inoculum was transferred to each well of the plate and mixed with 500 µl of liquified soft top agar (Nutrient Broth, NB, (Oxoid) with 7 g L^-1^ agar), precooled to 45 °C in a water bath. Plates were gently stirred (20 rpm) to ensure an even distribution of bacterial cells in the inoculated wells. After the agar had solidified, plates were left to dry for 10 min. in a laminar flow hood. Two µl of 10-fold diluted tailocins (approximately 10^10^ particles mL^-1^) purified from mitomycin-induced cultures of *D. dadantii* 3937 were spotted on the surface of the inoculated soft-top agar in the wells of the multitier plate. Inoculated plates were incubated for 24 h at 28 °C and then inspected for the presence of a growth inhibition. A spot of lack of growth was interpreted as the susceptibility of the given strain to the tailocins (positive reaction). Two µl of sterile PBS was spotted on a lawn of a suseptical strain as a negative control. The susceptibility of each bacterial strain was tested in triplicate, and the entire experiment was repeated two times.

### Determination of the bacterial killing rate

The rapidity by which tailocins killed bacteria was measured in 96-well plates (Nest) using a Epoch 2 microplate reader (BioTek). Twenty-five µl of a suspension containing tailocins (approximately 10^11^ particles mL^-1^) in PBS pH 7.2 were added to 100 µl of 5 McF (approximately 10^8^ CFU mL^-1^) bacterial suspension in PBS. An overnight culture of a tested strain in TSB were harvested by centrifugation (5 min, 8000 RCF). The OD_600_ of the PBS suspensions was measured each minute for 2 hours. The plate was incubated at 28 °C with shaking. The susceptible strain *M. paradisiaca* strain NCPPB 2511 was used as a positive control, and sterile PBS pH 7.2 without tailocins was used as a negative control. The log-transformed values of OD_600_ at each time point were normalized to the log-transformed starting OD_600_ and regressed against time. The regression coefficient (ΔLog_10_(OD_600_)min^-1^) was calculated for each of the obtained curves. The killing proportion was estimated at two representative time points (20 and 120 min) as the average % of the initial OD_600_ of the selected bacterial culture compared to controls (n=10).

### Analysis of the tailocins with SDS-PAGE and ESI LC-MS/MS

Proteins within tailocins were separated using a 4-20% sodium dodecyl sulfate-polyacrylamide gradient gel (Mini-PROTEAN TGX Stain-Free Precast, Bio-Rad Hercules, USA) electrophoresis (SDS-PAGE) using previous methods (*63*). Protein bands were excised from the gel using a sterile scalpel, and the excised gel pieces were placed in separate 1.5 mL Eppendorf tubes for amino acid sequencing. In-gel digestion was performed according to a standard protocol consisting of gel de-coloration and removal of Coomassie staining, reduction/alkylation with dithiothreitol (DTT), and iodoacetamide (IAA), respectively (*64*). First, digestion was carried out overnight with trypsin (Promega Mass Spectrometry Grade Gold) at 37 °C. The tryptic peptides were then eluted from the gel with sequential washing of gel pieces with 50 mM ammonium bicarbonate buffer, 5% formic acid in 50% acetonitrile, and 100% acetonitrile (*65*). All samples were then concentrated (SpeedVac), and the final cleanup was carried out using the StageTips method on the C18 phase to a 50% acetonitrile solution with 1% acetic acid (*66*). After concentrating the samples to 30 ml using the SpeedVac, fragmentation mass spectra were recorded for analysis. ESI LC-MS/MS analysis was performed on Triple Tof 5600+ mass spectrometer with DuoSpray Ion Source (AB SCIEX, Framingham, MA) connected to the Eksigent microLC (Ekspert MicroLC 200 Plus System, Eksigent, Redwood City, CA) equipped with the ChromXP C18CL column (3 μm, 120 Å, 150 × 0.5 mm). The microLC−MS/MS system was controlled by the AB SCIEX Analyst TF 1.6 software. Chromatographic separation was carried out for 30 min in a gradient program: (1) 0−1 min – 20% solvent B, (2) 1−26 min – 20−60% solvent B, (3) 26−28 min − 98% solvent B, (4) 29-30 min − 20% solvent B, where solvent A was 0.1% formic acid in water and solvent B 0.1% formic acid in acetonitrile. The identification of proteins present in the examined gel bands was carried out based on the obtained fragmentation spectra using the ProteinPilot software (v 4.5) or Peaks Studio and the appropriate protein database (UniProt *Dickeya dadanti 15.02.2023, unreviewed*) with an automated false discovery rate (1% FDR).

### Bioinformatic analyses

Prediction of prophage regions in the genome of *D. dadantii* 3937 (Genbank accession: NC_014500.1) was conducted using PHASTER (42). The tailocin cluster and individual proteins it encodes were analyzed against NCBI databases using BLASTn and BLASTp (43). The topology of the tailocin cluster in *D. dadantii* 3937 was investigated using BioCyc (44). Phylogenomic analysis of the Soft Rot *Pectobacteriaceae* (SRP) genomes was based on core genome sequences and was performed using EDGAR ver. 3.0 (45) accessed via https://edgar3.computational.bio.uni-giessen.de.

### Stability of tailocins (*43, 67, 68*)

Tailocins were incubated for 24 hours at several different temperatures (−80, −20, 4, 22, 28, 37, 42, 50, 65, and 80 °C), pH values (2, 5, 3.5, 7, 9, 10.5 and 12), and NaCl concentrations (0.01, 0.1, 1 and 5 M), after which their bacterial growth inhibitory activity was assessed by a spot test using *M. paradisiaca* as described above. All tested samples had a total volume of 200 µl, and the initial concentration of PEG-purified tailocin of approximately 10^10^ particles mL^-1^. To obtain the test samples, tailocins were diluted by mixing a volume of 20 µL of PEG-purified samples with 180 µl of either PBS (temperature stability), PBS with pH modified by addition of either HCl or NaOH to the desired values (pH stability), or in water containing different concentraitons of NaCl. The experiments were repeated three times, each with three technical replications per tested condition.

### Effect of enzyme/detergent/organic solvent treatment on the activity of tailocins (*67–69*)

Tailocins were incubated for 1 hour at 37 °C with the following enzymes: proteinase K (Sigma-Aldrich, final concentration: 0.5 mg mL^-1^), DNase (Sigma-Aldrich, final concentration: 10 U mL^-1^), or lysozyme (Sigma-Aldrich, final concentration: 0.5 mg mL^-1^), and at room temperature (22 °C) with sodium dodecyl sulfate (SDS, Sigma Aldrich, final concentration: 1%), or chloroform (POCH, 50% v/v). The remaining activity of treated tailocins was then assessed in a spot test as described above. The processed samples had a total volume of 200 µL, and a tailocin concentration of approximately 10^10^ particles mL^-1^. The samples were prepared by mixing 20 µl of PEG-purified samples (about 10^11^ particles mL^-1^) with the appropriate volume of the factor stock to a final volume of 200 µL with PBS. The exception were the chloroform-treated samples where 200 µL of sample were mixed on a shaker with 200 µL of chloroform. Prior to testing, the chloroform-treated sample was centrifuged (5 min at 4000 RCF), and the aqueous phase was removed for assay. The experiments were repeated three times, each with three technical repetitions per tested condition.

### Binding of tailocins to nonviable bacterial cells (*68*)

Overnight bacterial cultures of *M. paradisiaca* NCPPB 2511 and *D. dadantii* strain 3937 grown in TSB were washed twice with PBS buffer and then killed by incubation with 5 mg mL^-1^ chloramphenicol (Sigma-Aldrich) for 60 min, with shaking (120 rpm, 28 °C). The killing of the cells after 1 h was confirmed by plating 4 aliquots (10 μL) of treated culture on TSA plates, incubating at 28 °C for 24 h, and verifying last of bacterial growth. After killing, the nonviable bacterial cells were again washed three times with PBS to remove the remaining antibiotic. The PEG-purified tailocin samples were then added to suspension of viable and nonviable susceptible and nonsusceptible bacteria to a final concentration of approximately 10^10^ particles mL^-1^ and incubated for 40 min with shaking at 120 rpm at 28 °C. After incubation, samples were filtered through a 0.2 µm PES (polyether sulfone) membrane filter, and the remaining tailocins was assessed by a spot test using *M. paradisiaca*. The experiments were repeated three times, each with three technical replication per tested condition.

### Testing the influence of pH on the bactericidal activity of tailocins

The PEG-purified tailocins (approximately 10^11^ particles mL^-1^) were diluted 10-fold in PBS buffered to pH 2, 7, and 12 and incubated for 24 h at room temperature. The pH of each sample was then adjusted to neutral pH (pH 7) using an equal volume of the buffered PBS. The solutions were then incubated for 4 hours at room temperature and tested for growth inhibitory activity using a spot test in three replicates as described above.

#### Caenorhabditis elegans toxicity assay

Sensitivity of *Caenorhabditis elegans* to tailocins produced by *D. dadantii* may influence the was tested as before (*70*) with slight modifications. Briefly, wild-type Bristol N2 strain of *C. elegans* nematode obtained from the *Caenorhabditis* Genetic Center (CGC, University of Minnesota, Minneapolis, USA) was maintained as described before (*71, 72*) on Nematode Growth Medium (NGM) plates with *Escherichia coli* strain OP50 as a food source. The toxicity of the tailocins was tested on nematode cultures synchronized as described earlier (*73*). Briefly, nematode eggs were harvested from cultures treated with a mixture of 5M NaOH and 5.25% NaOCl (1:3, v:v) – a treatment that eliminates adult worms. Recovered eggs were hatched in an S-complete medium, and the nematodes were grown to the L4 stage using *E. coli* OP50 as a food source. The synchronized cultures were then transferred into wells of a 96-well plate. The worms were counted under the microscope (Leica MZ10f stereomicroscope, Leica) and ca. 30 worms placed in each well supplemented with PEG-purified and ultracentrifugation concentrated (1 h, 26 000 RCF, 4 °C) tailocin in an S-complete medium at a concentration of 10^12^ particles mL^-1^. Nematode cultures grown without tailocins were used as control. The plates were incubated for 24 h in the dark at 25 °C. The survival of nematodes in tailocin-supplemented cultures was compared to the control. The experiment was repeated twice with the same setup, and the results were averaged for analysis.

#### Statistical analysis

All statistical tests were conducted using either Past 4.13 software (*74*) or Microsoft Office Excel (www.office.com). The Shapiro-Wilk (*75*) and F-tests (*76*) were used to test for normality and variance equality of data, respectively. For pairwise testing, the t-test (*77*) was applied for data having a normal distribution and equal variances, the Welch test (*78*) was used for samples with a normal distribution but unequal variances, while the U Mann-Whitney test (*79*) was applied for data that were not normally distributed. One-way analysis of variance (ANOVA) (*80*) was used to compare more than two data groups. Levene’s test (*81*) was used to test the homogeneity of variance, and the normality of the residuals was conducted using the Shapiro-Wilk test. Welch’s one-way analysis of variance, followed by the Games-Howell post hoc test (*82*), was used for the data groups with non-homogeneous variance and normally distributed residuals. For groups with a non-homogeneous variance and without normally distributed residuals Kruscal-Wallis’s one-way analysis of variance (*83*) followed by Dunn’s post hoc test (*84*) was applied.

## Acknowledgments

The authors would like to express their gratitude to Dr. Nicole Hugouvieux-Cottee-Pattat (Laboratory of Microbiology Adaptation and Pathogenesis (UMR 5240) CNRS, Université Lyon 1 & INSA de Lyon, France) for providing strain A5587 (*D. dadantii* strain 3937 *tssK*::*uidA* kanR) for this study, Prof. Alfonso Jaramillo and Dr. Cristina Ramos (De Novo Synthetic Biology Lab, Institute for Integrative Systems Biology (I2SysBio-CSIC)) for providing valuable information on the P2 phage and Prof. Steven E. Lindow (University of California-Berkeley, Berkeley, CA, United States) for his comments on the manuscript and his editorial work.

## Author contributions

Conceptualization: RC, MB, DMK Methodology: MB, DMK, MN, MS, MR, PC, KW Investigation: MB, MS, MN, MR, DMK, KW Visualization: MB, DMK, MN, MR, Supervision: RC, DMK Writing—original draft: MB, DMK, RC Writing—review & editing: RC, DMK, MB Funding acquisition: RC

## Data and materials availability

All data are available in the main text or the supplementary materials.

