## Supplementary materials for "Soft rot pathogen *Dickeya dadantii* 3937 produces tailocins resembling the tails of *Enterobacteria* bacteriophage P2"

Marcin Borowicz *et al.*

**This PDF file includes:**

Figs. S1 to S4

Tables S1 to S2

References

**Other Supplementary Materials for this manuscript include the following:**

Data S1 to S2

### **Fig. S1**

### Host-dependent killing rate of P2D1 dickeyocins. The killing rate was calculated as the percentage (%) of the remaining viable bacterial cells of the susceptible *Dickeya* spp. strains (measured by OD_600_) after 20 min (A) and after 120 min (B) of their incubation with P2D1 dickeyocins. The results are shown as a box plot; whiskers reflect the maximum and minimum, box sides reflect the first and third quartile, and the bars reflect medians. Points indicate particular measurements (n=10). Statistically significant differences between the treatments were obtained using Welch's one-way analysis of variance followed by the Games-Howell post hoc test Groups with the same letter are not significantly different. (p<0.05).

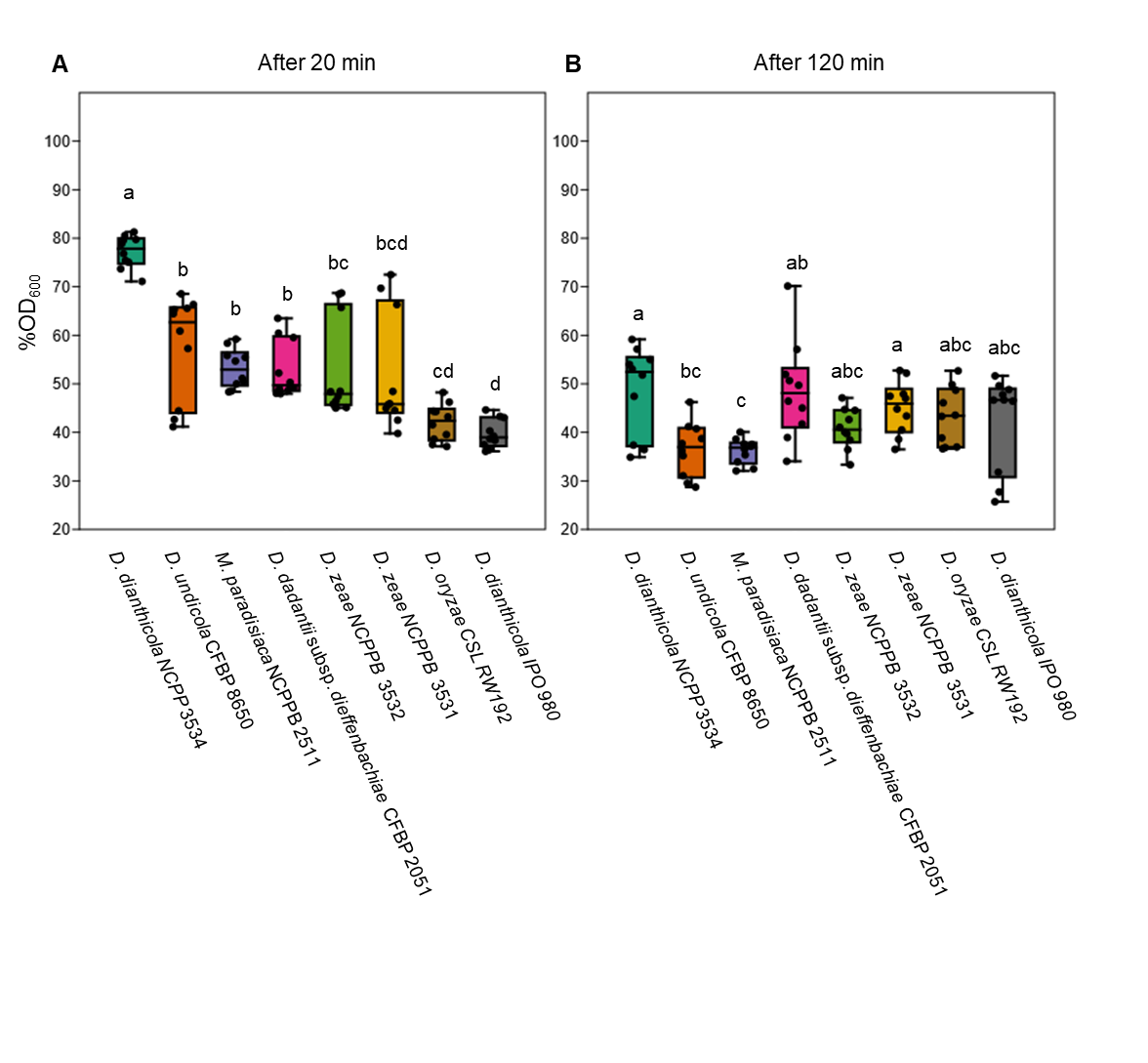

### **Fig. S2**

### Remaining activity of dickeyocins P2D1 after their binding to viable and nonviable (dead) cells of the susceptible (*M. paradisiaca* NCPPB 2511) and nonsusceptible (*D. dadantii* 3937) strains. Results are shown as box plots; the whiskers reflect the maximum and minimum, the box sides reflect the first and third quartile, bars reflect the medians. Points indicate particular measurements (n=9). Statistically significant differences between the treatment and the control were obtained using Kruscal-Wallis's one-way analysis of variance followed by Dunn's post hoc test. Groups with the same letter are not significantly different (p<0.05). AU – relative units.

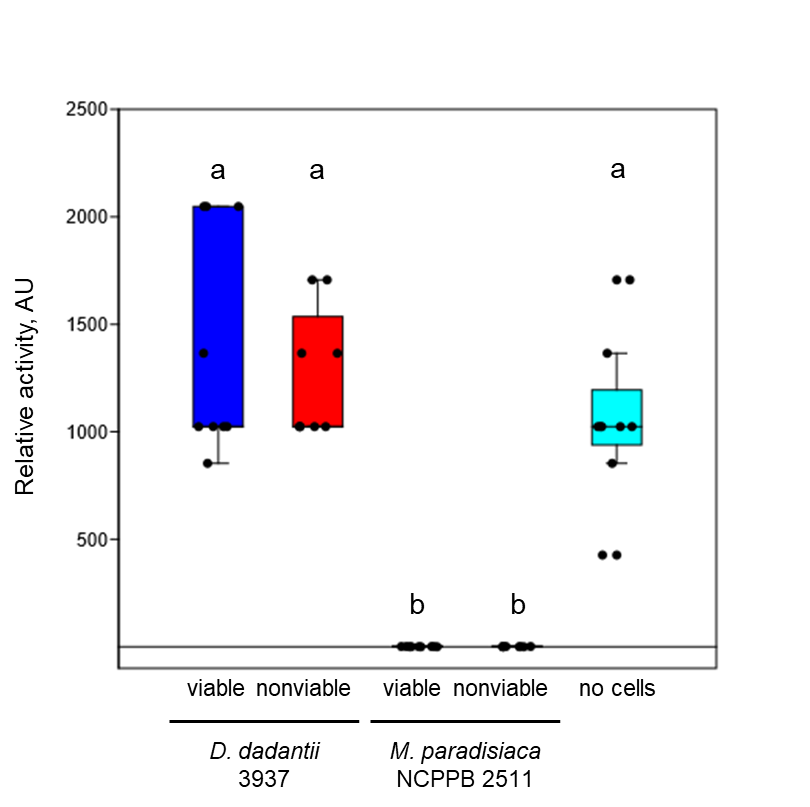

### **Fig. S3**

### Morphologic comparison of phage tail-like particles produced by T6SS defective mutant (A5587) and P2D1 dickeyocins produced by a wild type *D. dadanti* strain 3937. Representative TEM images of tailocins produced by the *Dickeya dadantii* wild-type strain and mutant A5587 (insertion in the *tssK* gene) are shown. The diameters (weight and length) were compared using TEM, with *n* representing the number of particles measured to obtain the average.

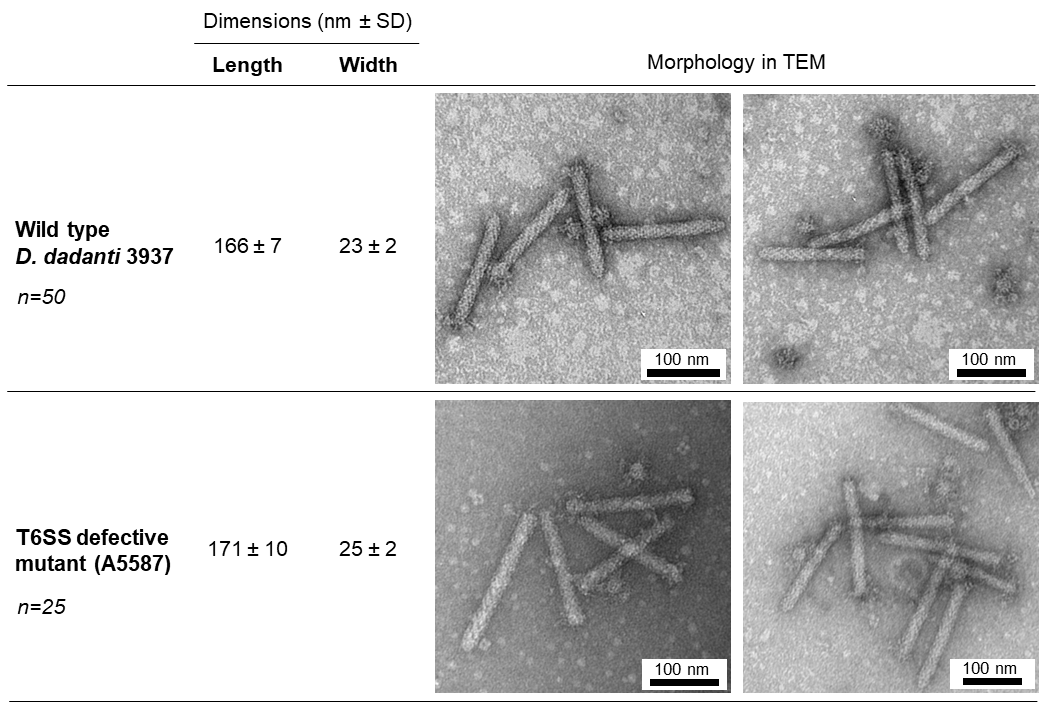

### **Fig. S4**

### Survival of *C. elegans* in the presence of P2D1 dickeyocins in the liquid killing assay. *C. elegans* growth medium supplemented with PBS without tailocins served as a control. Eight different concentrations of P2D1 were tested. Results are shown as box plots; the whiskers reflect the maximum and minimum, the box sides reflect the first and third quartile, bars reflect the medians. Points indicate particular measurements (n=9).

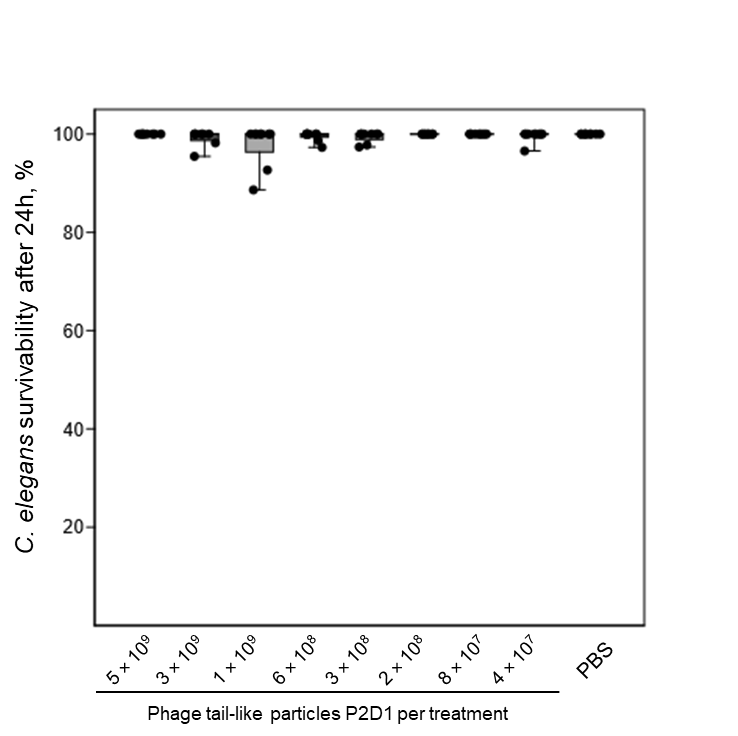

### **Table S1**

### List of strains used in this study.

| **Strain** | **Host plant/origin of isolation** | **Geographical origin, year of isolation** | **Other collection numbers** | **Ref.** |
| --- | --- | --- | --- | --- |
| **Soft Rot *Pectobacteriaceae*** | | | | |
| *Dickeya chrysanthemi* NCPPB 3533 | *Solanum tuberosum* | United States, 1987 | IFB0139 | *(1)* |
| *Dickeya chrysanthemi* NCPPB 402 | *Chrysanthemum morifolium* | United States, 1956^*^ | IFB0055, CFBP 2048, ATCC 11663 | *(1, 2)* |
| *Dickeya chrysanthemi* NCPPB 516 | *Parthenium argentatum* | Denmark, 1957^*^ | IFB0724, CFBP 1270 | *(1)* |
| *Dickeya dadantii* 3937 | *Saintpaulia sp.* | France, 1972 | IFB0016 | *(3)* |
| *Dickeya dadantii* DSM 18020 | *Pelargonium capitatum* | Comoros, 1961^*^ | IFB0010, CFBP 1269, NCPPB 898, SCRI 1269 | *(2)* |
| *Dickeya dadantii* NCPPB 3537 | *Solanum tuberosum* | Peru, 1987 | IFB0127 | *(1)* |
| *Dickeya dadantii* subsp. *dieffenbachiae* NCPPB 2976 | *Dieffenbachia* | United States, 1977^*^ | IFB0718, CFBP 2051 | *(1, 2)* |
| *Dickeya dianthicola* IPO 980 | *Solanum tuberosum* | Netherlands | IFB0140 | *(4)* |
| *Dickeya dianthicola* NCPPB 3534 | *Solanum tuberosum* | Netherlands, 1987 | IFB0126 | *(4)* |
| *Dickeya fangzhongdai* DSM 101947 | *pear tree (bleeding cancer)* | China, 2009 | IFB0716, CFBP 8607 | *(5)* |
| *Dickeya lacustris* CFBP 8647 | *water* | France, 2017 | IFB0715, LMG 30899 | *(6)* |
| *Dickeya oryzae* CSL RW 192 | *river water* | England | IFB0220 | *(1)* |
| *Dickeya poaceiphila* NCPPB 569 | *Saccharum officinarum* | Australia, 1958 | IFB0717, CFBP 8731 | *(7)* |
| *Dickeya solani* D s0432-1 | *Solanum tuberosum* | Finland, 2004 | IFB0135, IPO 3295, LMG 27551 | *(8)* |
| *Dickeya solani* GBBC 2040 | *Solanum tuberosum* | Belgium, 2007^*^ | IFB0484, LMG 25865 | *(4)* |
| *Dickeya solani* MK10 | *Solanum tuberosum* | Israel | IFB0723 | *(4)* |
| *Dickeya solani* MK16 | *river water* | United Kingdom | IFB0272, IPO 3494 | *(1, 4)* |
| *Dickeya* sp. CSL RW 240 | *river water* | England | IFB0721 | *(1)* |
| *Dickeya* sp. MK7 | *river water* | Scotland | IFB0275 | *(1)* |
| *Dickeya* sp. NCPPB 3274 | *Aglaonema* | St. Lucia, 1983 | IFB0722 | *(1)* |
| *Dickeya undicola* CFBP 8650 | *water* | Malaysia, 2014 | IFB0714, LMG 30903 | *(9)* |
| *Dickeya zeae* MK19 | *river water* | Scotland | IFB0719 | *(1)* |
| *Dickeya zeae* NCPPB 3531 | *Solanum tuberosum* | Australia, 1987^*^ | IFB0138 | *(1)* |
| *Dickeya zeae* NCPPB 3532 | *Solanum tuberosum* | Australia, 1987^*^ | IFB0720 | *(1)* |
| *Musicola paradisiaca* NCPPB 2511 | *Musa paradisiaca* | Colombia, 1973^*^ | IFB0117, ATCC 33242, LMG 2542 | *(1, 10)* |
| *Pectobacterium actinidiae* LMG 26003 | *Actinidia chinensis* | Korea | IFB5641 | *(11)* |
| *Pectobacterium aroidearum* NCPPB 929 | *Zantedeschia aethiopica* | South Africa, 1959 | IFB5514, LMG 2417 | *(12)* |
| *Pectobacterium atrosepticum* NCPPB 549 | *Solanum tuberosum* | United Kingdom, 1957 | IFB5399, CFBP1526, ATCC 33260 | *(13)* |
| *Pectobacterium atrosepticum* SCRI 1043 | *Solanum tuberosum* | Scotland, 1985 | IFB5102 | *(14)* |
| *Pectobacterium betavasculorum* CFBP 2122 | *Beta vulgaris cv. Saccharata* | USA, 1971 | IFB5269, CFBP 2122, NCPPB 2795 | *(13)* |
| *Pectobacterium brasiliense* LMG 21371 | *Solanum tuberosum* | Brazil, 1999 | IFB5390, ATCC BAA-417 | *(15)* |
| *Pectobacterium cacticida* CFBP 3628 | *Carnegiea gigantea* | USA, 1944 | IFB5644, ATCC 49481, CIP 105191 | *(16)* |
| *Pectobacterium carotovorum* CFBP 2046 | *Solanum tuberosum* | Denmark, 1952 | IFB5263, NCPPB 312, ATCC 15713 | *(16)* |
| *Pectobacterium fontis* CFBP 8629 | *water* | Malaysia, 2015 | IFB5645, LMG30744 | *(17)* |
| *Pectobacterium parmentieri* CFBP 8475 | *Solanum tuberosum* | France, 2008 | IFB5648, LMG 29774 | *(18)* |
| *Pectobacterium parmentieri* SCC3193 | *Solanum tuberosum* | Finland, 1980s | IFB5395 | *(19)* |
| *Pectobacterium peruviense* CFBP 5834 | *Solanum tuberosum* | Peru, 1979 | IFB5232, LMG 30269; PCM 2893; SCRI 179 | *(20)* |
| *Pectobacterium polaris* NCPPB 4611 | *Solanum tuberosum* | Norway, 2010 | IFB5646, CFBP 8603 | *(21)* |
| *Pectobacterium polonicum* DPMP 315 | *vegetable field* | Poland, 2016 | IFB5673, LMG 31077 | *(22)* |
| *Pectobacterium punjabense* CFBP 8604 | *Solanum tuberosum* | Pakistan, 2017 | IFB5642, LMG30622 | *(23)* |
| *Pectobacterium versatile* CFBP 6051 | *Solanum tuberosum* | Netherlands, 2001 | IFB5636, NCPPB 3387 | *(15)* |
| **Other strains** | | | | |
| *Citrobacter freundii* ATCC 8090 |  | Unknown, 1928 | NCTC 9750 | *(24)* |
| *Escherichia coli* ATCC 25922 | *clinical isolate* | USA, 1946 | DSM 1103, NCIB 12210 | *(25)* |
| *Escherichia coli* ATCC 8739 | *feces* |  |  | *(26)* |
| *Escherichia coli* OP50 |  |  |  | *(27)* |
| *Klebsiella aerogenes* ATCC 51697 |  |  |  | *(28)* |
| *Klebsiella quasipneumoniae* ATCC 700603 |  |  | K6, CCUG 45421, LMG 20218 | *(29)* |
| *Pseudomonas aeruginosa* PA14 | *clinical islate* |  | DSM 19882 | *(30)* |
| *Pseudomonas aeruginosa* PAO1 | *clincal isolate* | Australia, 1954 | DSM 22644, ATCC 15692 | *(31)* |
| *Pseudomonas donghuensis* P482 | *Solanum lycopersicum* | Poland, 2012 |  | *(32)* |
| *Serratia marcescens* ATCC 14756 |  | USA | PCI 1107 | *(33)* |
| *Staphylococcus aureus* ATCC 25923 | *clinical isolate* |  |  | *(34)* |

* – the year of addition to the collection

NCPPB – National Collection of Plant Pathogenic Bacteria

CFBP – French Collection for Plant Associated Bacteria

DSM – German Collection of Microorganisms and Cell Cultures GmbH

ATCC – American Type Culture Collection

LMG (BCCM) – Belgian Coordinated Collections of Micro-organisms

### **Table S2**

### Determination of the concentration of P2D1 dickeyocins induced from *D. dadantii* strain 3937. To determine the concentration of the dickeyocins, three independent methods were used: (i) semiquantitative estimation by a spot test, (ii) Poisson distribution killing method, and (iii) direct particle count with NanoSight NS300. For the estimation of the number of dickeyocins, three independent inductions of P2D1 dickeyocins (=3 biological replicates) were done. Likewise, the induction of phage tail-like particles from T6SS defective mutant of *D. dadantii* strain 3937 was done using the same protocol as used for P2D1 dickeyocins and described in the Materials and Methods section. The results are shown as relative units (AU) mL^-1^ ± standard deviation, killing particles mL^-1^ ± standard deviation, and particles mL^-1^ ± standard error, respectively.

| Sample | | Concentration determination method | | |
| --- | --- | --- | --- | --- |
|  |  | **Semiquantitative estimation** (spot test, AU mL^-1^  ± standard deviation) | **Poisson distribution killing method,**  (killing particles mL^-1^  ± standard deviation) | **Direct particle count with NanoSight NS300**,  (particles mL^-1^  ± standard error) |
| Inductions from strain 3937 | I | 2.2±1.0 × **10^6^** | 1.6±1.3 × **10^11^** | 2.8±0.6 × **10^10^** |
|  | II | 7.2±5.1 × **10^6^** | 2.7±1.9 × **10^11^** | 1.5±0.2 × **10^11^** |
|  | III | 4.8±1.7 × **10^6^** | 2.9±1.7 × **10^11^** | 8.8±0.5 × **10^10^** |
| Induction from the T6SS defective 3937 mutant | | 3.8±0.8 × **10^6^** | 2.1±1.4 × **10^11^** | 8.2±1.0 × **10^10^** |
| PEG purification followed by ultracentrifugation | | 6.1±2.2 × **10^6^** | 1.2±1.1 × **10^12^** | 1.0±0.1 × **10^12^** |

Data S1 (separate file)

NCBI database searches using blastp and blastn.

Data S2 (separate file)

Comparison of sequence homology between proteins of P2D1 and phage P2.
